# A balance between nucleating and elongating actin filaments controls deformation of protein condensates

**DOI:** 10.1101/2025.06.18.660423

**Authors:** Caleb Walker, Daniel Mansour, Unyime Effiong, Dominique Jordan, Liping Wang, Eileen M. Lafer, Jose Alvarado, Brian Belardi, Padmini Rangamani, Jeanne C. Stachowiak

**Affiliations:** Department of Biomedical Engineering, The University of Texas at Austin, Austin, TX, USA; Department of Mechanical and Aerospace Engineering, University of California, San Diego, La Jolla, CA, USA; McKetta Department of Chemical Engineering, The University of Texas at Austin, Austin, TX, USA; Department of Biochemistry and Structural Biology, University of Texas Health Science Center at San Antonio, San Antonio, TX, USA; Department of Physics, The University of Texas at Austin, Austin, TX, USA; Department of Pharmacology, University of California, San Diego School of Medicine, La Jolla, CA, USA

## Abstract

Protein condensates use multivalent binding and surface tension to assemble actin filaments into diverse architectures, reminiscent of filopodia and stress fibers. During this process, nucleation of new filaments and elongation of existing filaments inherently compete for a shared pool of actin monomers. Here we show that a balance between these competing processes is required to deform condensates of VASP, an actin binding protein, into structures with high aspect ratios. Addition of magnesium, which promotes filament nucleation, helped actin to deform condensates into high aspect ratio structures. In contrast, addition of profilin, which inhibits filament nucleation, allowing existing filaments to elongate, caused actin to assemble into ring-like bundles that failed to substantially increase condensate aspect ratio. Computational modeling helped to explain these results by predicting that a group of short linear filaments, which can apply asymmetric pressure to the condensate boundary, is needed to increase condensate aspect ratio. In contrast, a small number of long filaments with the same total actin content should fail to overcome the droplet surface tension, forming a ring-like bundle. To test these predictions, we introduced gelsolin, which severed log filaments within rings, creating new barbed ends. The resulting set of shorter filaments regained the ability to deform condensates into high aspect ratio structures. Collectively, these results suggest that a balance of actin filament nucleation and elongation is required to deform protein condensates. More broadly, these findings illustrate how protein condensates can balance multiple kinetic processes to direct the assembly of diverse cytoskeletal architectures.

## Introduction

The actin cytoskeleton facilitates critical cellular processes such as motility, endocytosis, morphogenesis, and intracellular transport.^1–4^ Actin filaments (F-actin) are assembled from monomeric (G-actin) subunits through tightly regulated polymerization processes that are orchestrated by a diverse array of actin-binding proteins.^3,5–9^ To maintain the structural and functional plasticity of actin networks, these regulatory factors govern every stage of filament assembly, including nucleation, elongation, severing, capping, and bundling.^3,5–7^ Recently, several actin accessory proteins have been shown to undergo liquid–liquid phase separation (LLPS), a process by which biomolecules self-associate into a dense, liquid-like phase surrounded by a dilute phase.^10–12^ Initial studies showed that multivalent interactions among signaling proteins such as Nck, N-WASP, and LAT, and can drive the formation of liquid-like condensates, and that these condensates locally concentrate actin nucleation factors, thereby enhancing assembly of actin filaments.^13,14^ Subsequent work showed that actin assembly can be modulated by adjusting protein stoichiometry, highlighting how compositional changes within condensates directly regulate actin polymerization.^15^ Recent work has expanded the scope of phase separation in cytoskeletal regulation to include proteins that control microtubule assembly, such as SPD-5 and γ-tubulin within centrosomal condensates,^16^ and other actin accessory proteins like abLIM1.^17^ Building on this framework, recent experiments have shown that condensates composed of the actin polymerase and bundling protein, VASP, facilitate the assembly of actin filaments, which cause the condensate to progress through a series of morphological changes that end in deformation into rod-like structures filled with a bundle of linear actin filaments.^18^ This transition is driven by the interplay between filament rigidity and the surface tension of the condensate, resulting in the accumulation of actin filaments at the inner surface of spherical condensates, followed by the assembly of filaments into ring-like bundles.^19^ With the addition of more actin filaments, these bundles eventually overcome the condensate surface tension, deforming the initially spherical condensate into a linear, rod-like morphology filled with parallel actin filaments.^18^ Further work showed that the addition of Arp2/3 to these VASP condensates facilitated branched actin networks, leading to multiple protrusions from the droplet.^20^ Complementary computational studies demonstrated that these morphological outcomes reflect kinetic constraints on filament bundling and rearrangement.^19^ Extending this framework, we demonstrated that multivalent condensates composed of actin-binding proteins can drive both filament nucleation and elongation, even in the absence of dedicated polymerase or nucleation proteins. These findings showed that condensate-mediated actin assembly can result from multivalent interactions with actin, likely promoting nucleation and elongation of nascent filaments.^21^

Although many actin-binding proteins are known to regulate actin assembly in cells, how these regulatory inputs influence condensate-mediated actin assembly and final condensate morphology remains unclear. In particular, when the supply of monomeric actin is limited, as is the case for local actin assembly in cells or in reconstituted systems, there is an inherent competition between the nucleation of new filaments and the elongation of existing ones. These two processes draw from the same pool of actin monomers, such that conditions that favor nucleation would result in many shorter filaments, while those that favor elongation would produce fewer but longer filaments. This tradeoff has recently been demonstrated in competition between formin-mediated nucleation and elongation processes,^22^ and also in competition between Arp2/3 mediated nucleation and formin-driven nucleation and elongation.^23^

We reasoned that, within condensates, where actin-binding and regulatory proteins can be locally concentrated, this tradeoff may be amplified, potentially influencing the resulting actin network and condensate architecture. Therefore, we asked how altering the kinetics of filament assembly, through divalent cation exchange or addition of regulatory proteins, would affect condensate architecture. Magnesium promotes filament nucleation by lowering the critical concentration for assembly.^24–26^ Conversely, profilin is known to sterically inhibit filament nucleation but can facilitate enhanced barbed-end elongation in the presence of elongation factors such as VASP.^9,27–30^ By tuning the balance between nucleation and elongation, we asked whether these regulators could shift the final condensate morphology. We found that exchanging Ca²⁺ for Mg²⁺, the physiologically relevant divalent cation, enhanced condensate-mediated filament assembly and condensate deformation. We then examined how the addition of profilin, resulting in profilin-actin, a more physiologically relevant monomer pool,^5,31^ altered the assembly of actin networks by VASP condensates. These experiments revealed that profilin suppresses the deformation of condensates into high-aspect-ratio structures, leading to dense, ring-like bundles of actin filaments trapped within condensates. Guided by agent-based simulations, we asked whether introducing filament turnover could release trapped filaments from toroidal arrangements. When we introduced the filament-severing protein gelsolin, toroids disappeared, and condensates were once again deformed into rod-like structures by actin assembly. These results provide insight into the balance between filament nucleation and elongation during actin-mediated deformation of protein condensates.

## Results and Discussion

### Magnesium enhances condensate-mediated assembly of actin filaments and bundles

A central factor in filament assembly dynamics is the identity of the bound divalent cation.^24,25,32^ In vitro, actin is often purified in the calcium-bound form, yet within cells, magnesium-bound actin is the physiological species.^5,25^ Mg^2+^-actin is also known to assemble more rapidly, with a lower critical concentration for filament formation than Ca^2+^-actin.^25,26,32^ Previous work in our lab demonstrated that VASP condensates mediate actin assembly using Ca²⁺-actin.^18,20^ However, the effect of directly exchanging Ca²⁺ for Mg²⁺ on actin filament assembly within VASP condensates has not been explored. To better reflect cellular conditions of actin filament assembly in vitro, we first exchanged actin from calcium to magnesium through the inclusion of 1 mM MgCl2 and 1 mM EGTA.^28,33^ As EGTA has a much stronger affinity for Ca^2+^ than Mg^2+^, it facilitates the sequestration of Ca^2+^ and subsequent exchange of actin to Mg^2+^. To explore the effect of magnesium on condensate-mediated filament assembly, we added 3 µM actin to VASP condensates formed in experimental buffer (20 mM Tris pH 7.4, 150 mM NaCl, 5 mM TCEP, 200 µM ATP). (**Fig. 1A)** Upon the addition of Ca^2+^-actin, we saw the characteristic sphere-to-rod transition **(Fig. 1B)** that has been described in our previous work.^18,20^ Briefly, as actin filaments assembled inside condensates and elongated, the condensates were transformed from their initial spherical morphology to a rod-like morphology. Upon exchange to Mg^2+^-actin, the fraction of condensates that underwent this transformation, defined as having an aspect ratio greater than 1.2, remained similar to that with Ca^2+^-actin **(Fig. 1C)**. However, there was a significant increase in the average aspect ratio of the condensates **(Fig. 1D,E)**, suggesting that the exchange from Ca^2+^-actin to Mg^2+^-actin enhanced filament assembly. To investigate this effect, we performed a pyrene-actin assay to monitor filament assembly kinetics.^34^ G-actin, containing 10% pyrene-labeled actin, was mixed with freshly prepared VASP condensates, formed by 3% PEG addition, with or without MgCl₂ and EGTA in the buffer. After combining the actin and condensates, pyrene fluorescence was monitored using a plate reader to quantify the bulk filament assembly kinetics. Consistent with our morphological data, Mg²⁺ exchange resulted in a marked increase in the rate of condensate-mediated filament assembly **(Fig. 1F).** Full pyrene assay curves are presented in **Fig. S2**.

**Figure 1.**
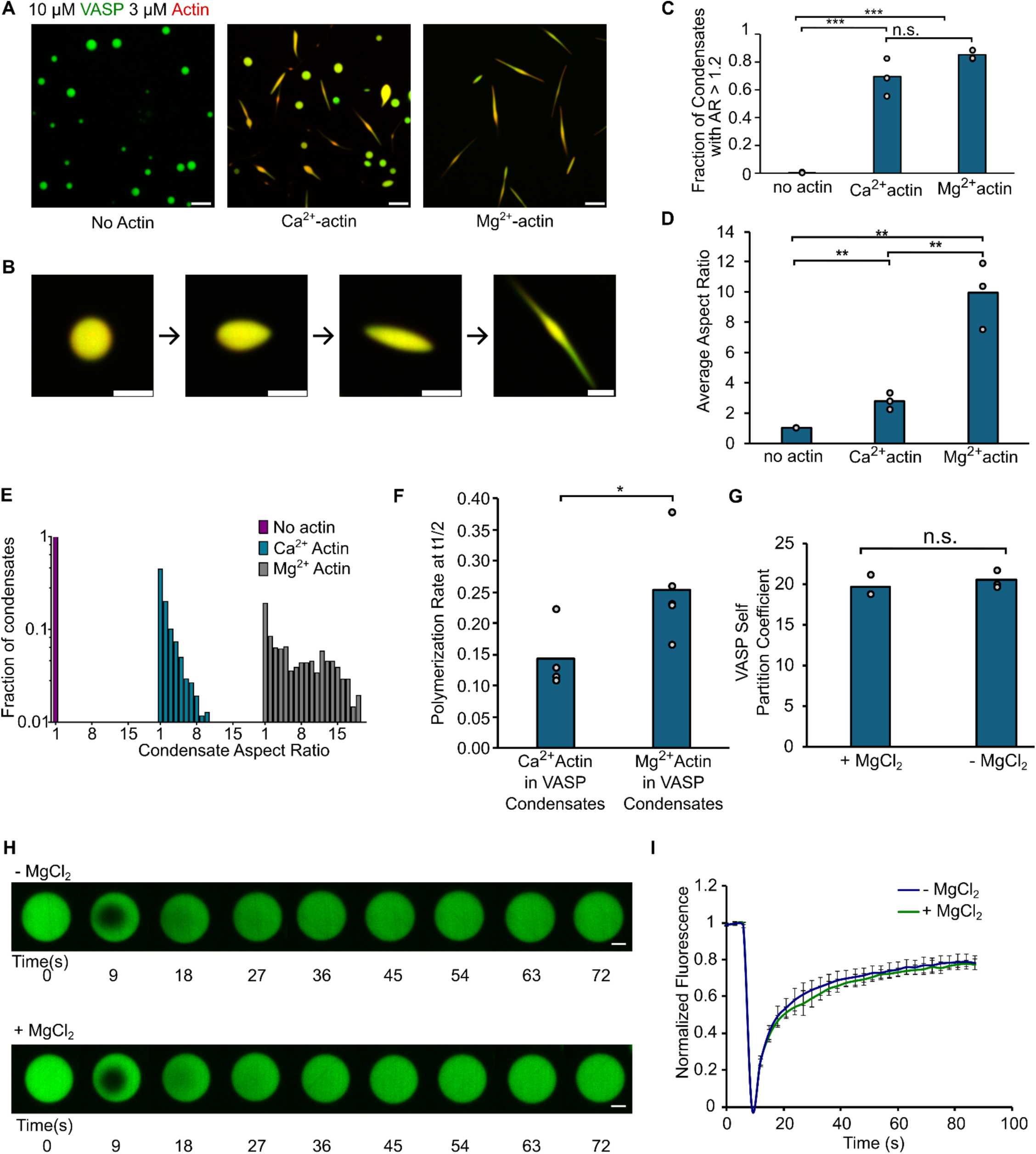
Magnesium enhances condensate-mediated assembly of actin filaments and bundles. A) Condensates formed from 10 µM VASP with the addition of 3 µM actin are increasingly deformed with the inclusion of MgCl2 and EGTA in the buffer to facilitate exchange to Mg^2+^-actin. Scale bars 5 μM. B) Representative images depicting the progression of condensate deformation from actin filament assembly. C) Quantification of the fraction of condensates with deformed morphologies, defined as having an aspect ratio greater than 1.2, for the conditions shown in A. Data are the mean across three independent experiments with at least 700 condensates analyzed per condition. Overlaid gray circles denote the means of each replicate. Three asterisks denote p<.001 using an unpaired, two-tailed t test on the means of the individual replicates, n=3. D) Quantification of the average condensate aspect ratio for the conditions shown in A. Data are means across three independent experiments with at least 700 condensates analyzed per condition. Overlaid gray circles denote the means of each of the replicates. Two asterisks denote p<.01 using an unpaired, two-tailed t test on the means of the individual replicates, n=3. E) Histograms showing the distribution of condensate aspect ratios for the conditions shown in A. For the Mg^2+^actin condition, condensates with aspect ratios greater than 20, corresponding to 9.7% of the data, were hidden to better visualize the distributions across all conditions. F) Quantification of filament assembly rates at t1/2 for pyrene actin assays for VASP condensates with Ca^2+^-actin and Mg^2+^-actin. Overlaid gray circles denote the means of each replicate. One asterisk denotes p<.05 using an unpaired, two-tailed t test on the calculated t1/2 values from the individual replicates, n=3. G) Self partitioning of 10 µM VASP into condensates formed in standard experimental buffer (20 mM Tris pH 7.4, 150 mM NaCl, 5 mM TCEP, and 200 µM ATP) with or without 1 mM EGTA and 1 mM MgCl₂ added. Data are the mean across three independent experiments with at least 450 condensates analyzed per condition. Overlaid gray circles denote the means of each replicate. No significance was found using an unpaired, two-tailed t-test on the means of the individual replicates, n=3. H) Representative images of fluorescent recovery after photobleaching of VASP condensates in buffer conditions with and without added MgCl2 and EGTA. Scale bars 2 µm. I) Plot of average fluorescence recovery +- SD after photobleaching for VASP condensates across n = 10 independent samples for buffer conditions with and without added MgCl2 and EGTA.

Because filament morphology within condensates arises from the interplay between filament rigidity and condensate surface tension, we also investigated whether the observed morphological differences could be attributed to changes in the condensates themselves. For this purpose, we formed VASP condensates with and without the addition of 1 mM EGTA and 1 mM MgCl2. We found no significant effect on either the self-partitioning of VASP condensates (**Fig. 1G)** or on the fluorescence recovery after photobleaching of VASP condensates (**Fig. 1H,I)**, suggesting that increased condensate deformation by Mg^2+^-actin was most likely the result of changes in the kinetics of actin assembly rather than changes in the physical properties of VASP condensates.

### Profilin limits condensate deformation without inhibiting actin filament assembly

Having observed that the exchange of actin from Ca^2+^ to Mg^2+^ enhanced actin-mediated condensate deformation, we reasoned that other actin regulatory factors known to modulate filament assembly kinetics might also influence these morphological outcomes. Profilin, a key regulator of monomer availability, sequesters actin and inhibits spontaneous nucleation, thereby controlling the spatial and temporal dynamics of filament growth.^9,27,35^ Additionally, profilin can accelerate barbed-end elongation in the presence of polymerases like VASP by delivering actin monomers via its polyproline-binding interface.^28,29,36^ Given its dual role in suppressing nucleation and promoting elongation, and as most cytosolic actin is bound to profilin in cells,^9,31^ we sought to determine how profilin-actin influences condensate-mediated filament assembly.

We first tested whether profilin affects the self-assembly and liquid-like nature of VASP condensates. Despite profilin’s strong partitioning into VASP condensates (**Fig. 2A)**, with profilin being enriched about 18-fold in the condensate phase, increasing profilin concentrations had no significant effect on VASP self-partitioning **(Fig. 2B)**. Additionally, VASP condensates displayed similar fluorescent recovery after photobleaching with increasing concentrations of profilin (**Fig. S1).** These observations suggest that profilin does not substantially disrupt VASP-VASP interactions or alter multivalent interactions required for VASP condensation. We next evaluated the impact of profilin on condensate-mediated actin assembly. Interestingly, we found that incubating actin with increasing concentrations of profilin before addition to VASP condensates resulted in a decrease in condensate deformation **(Fig. 2C)**, as measured by decreases in the fraction of condensates that exceeded a threshold aspect ratio of 1.2 (**Fig. 2D),** average aspect ratio of deformed condensates (**Fig. 2E),** and distribution of condensate aspect ratios **(Fig. 2F).** To test whether profilin inhibited condensate-mediated filament assembly, we used phalloidin to stain specifically for polymerized actin. Even with 15 µM profilin (corresponding to a 5:1 profilin:actin ratio), we observed rings of polymerized actin within the VASP condensates **(Fig. 2G),** suggesting that actin filament assembly continued to occur, despite the reduction in condensate deformation. However, pyrene-actin assays showed that profilin significantly reduced the filament assembly rate, consistent with its known role in suppressing spontaneous nucleation (**Fig. 2H)**. These results suggest that profilin partially inhibits condensate-mediated actin assembly, likely by limiting nucleation of new filaments. As a result, filaments appear to become trapped in ring-like architectures, failing to undergo the symmetry-breaking transitions required to elongate condensates into rod-like structures **(Fig. 2I)**.

**Figure 2:**
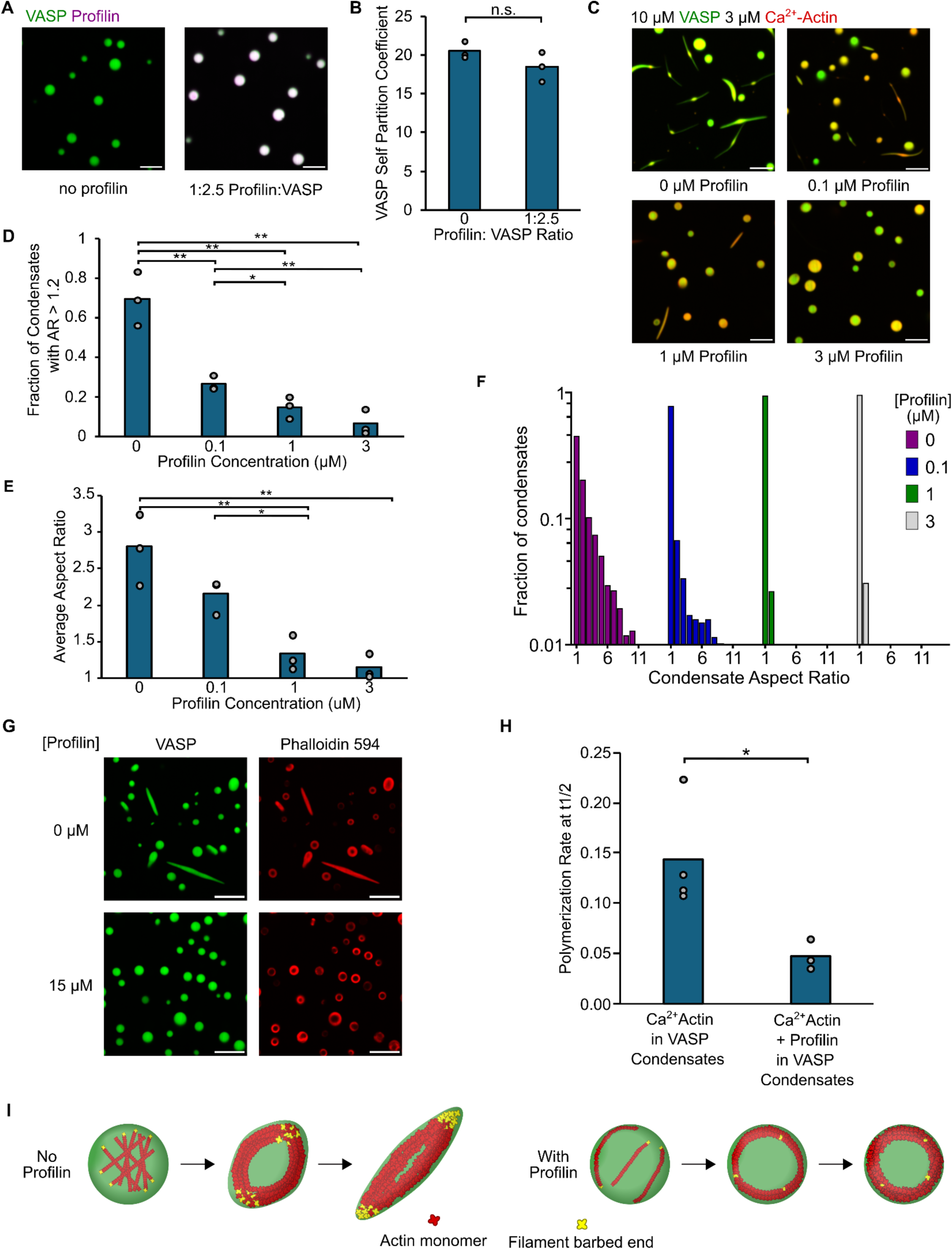
Profilin limits condensate deformation without inhibiting actin filament assembly. A) Representative confocal fluorescence images of VASP condensates (10 µM) formed in the absence and presence of Alexa Fluor 647- labeled profilin. Scale bars 5 μm. B) Quantification of the self-partitioning of VASP into condensates for the conditions shown in A. Data are the mean across three independent experiments, with the overlaid gray circles denoting the mean of each replicate. There is no statistically significant difference for any conditions using a two-tailed t-test on the means of the individual replicates, n=3. C) Incubation of 3 µM actin (red) with increasing concentrations of profilin (unlabeled) prior to addition to 10 µM VASP (green) condensates results in a decrease in condensate deformation. Scale bars 5 μm. D) Quantification of the fraction of condensates with deformed morphologies, defined as having an aspect ratio greater than 1.2, for the conditions shown in C. Data are means across three independent experiments with at least 1400 condensates analyzed per condition. Overlaid gray circles denote the means of each replicate. One asterisk denotes p<0.05, and two asterisks denote p<0.01 using an unpaired, two-tailed t test on the means of the individual replicates, n=3. E) Quantification of the average condensate aspect ratio for the conditions shown in C. Data are means across three independent experiments with at least 1400 condensates analyzed per condition. Overlaid gray circles denote the means of each replicate. One asterisk denotes p<0.05, and two asterisks denote p<0.01 using an unpaired, two-tailed t test on the means of the individual replicates, n=3. F) Histograms showing the distribution of condensate aspect ratios for the conditions shown in C. Condensates with aspect ratios greater than 15 (corresponding to 0.4%, 1.1%, 0.27%, and 0.28% of the data respectively for the 0, 0.1, 1, and 3 µM conditions) were hidden to better visualize the distributions across all conditions. G) Phalloidin-iFluor-594 (red) staining of VASP condensates (green) with 3 μM monomeric G-actin (unlabeled), and either no profilin or 15 μM profilin (unlabeled), displaying rings of filamentous actin within the protein condensates even in the undeformed conditions. Scale bars, 5 μm. H) Quantification of filament assembly rates at t1/2 for pyrene actin assays for VASP condensates and Ca^2+^-actin with and without 3 μM profilin included. Overlaid gray circles denote the means of each replicate. One asterisk denotes p<.05 using an unpaired, two-tailed t test on the calculated t1/2 values from the individual replicates, n=3. I) Cartoon depicting the proposed method that leads to inhibited condensate deformation with profilin present. (Left) With no profilin, actin assembly leads to condensate deformation. (Right) With profilin present, actin assembly is inhibited, making filaments unable to deform condensates. Buffer conditions for all experiments were (20 mM Tris pH 7.4, 150 mM NaCl, 5 mM TCEP, 200 μM ATP)

### The combination of magnesium and profilin traps actin filaments and condensates in toroidal morphologies

Next we investigated the impact of adding both magnesium and profilin simultaneously, mimicking the in vivo setting. In the presence of magnesium, the addition of a relatively low concentration of profilin (0.1 µM, 1:30 profilin:actin ratio) resulted in a modest decrease in condensate deformation, reflected by slightly lower average aspect ratios, but the condensates remained predominantly rod-like. As profilin concentration increased to 1 µM (1:3 profilin:actin ratio), deformation was further reduced. Surprisingly, a distinct toroidal morphology began to emerge under these conditions. This toroid structure became the dominant morphology at 3 µM profilin (a 1:1 profilin:actin ratio), replacing rod-like shapes as the primary condensate phenotype **(Fig. 3A).** Here we define toroids as ring-shaped structures with a visible central void in the condensate phase in both cross-section and 3D morphology, in contrast to the rings of actin within condensates, where the protein condensate is a continuous sphere or disc. This toroidal transition with increasing profilin was also characterized by a decrease in the fraction of condensates that displayed higher aspect ratios **(Fig. 3B),** a decrease in the average aspect ratio of the condensates **(Fig. 3C)**, and a shift in the distribution of condensate aspect ratios toward smaller values (**Fig. 3D).** Pyrene-actin assays using 3 µM profilin and Mg²⁺-actin showed a reduction in filament assembly rate compared to Mg²⁺-actin, as expected. However, the assembly rate remained higher than that observed for VASP condensates with Ca²⁺-actin in the absence of profilin **(Fig. 3E).** The fraction of condensates that displayed a toroidal morphology increased significantly with increasing profilin concentration **(Fig. 3F)**. These results suggest that although Mg²⁺ enhances overall filament assembly, the inclusion of profilin tempers this effect, limiting nucleation or altering filament growth dynamics, in a way that still permits actin filament assembly but changes the morphological outcome. Rather than promoting deformation of condensates into rod-like shapes, the combination of profilin and magnesium appears to stabilize toroidal architectures **(Fig. 3G-J).** The assembly of toroids, with a void in the center of the condensate, suggests a ring of actin that grew sufficiently rigid that it could no longer be contained in a spherical condensate. As we have reported previously,^18^ expansion of ring-like actin bundles can flatten spherical condensates into disc-like morphologies. Toroids suggest the next step in this progression, where ring-like actin bundles become more and more rigid, increasing in diameter until the limited volume of the condensate can no longer span the center of the ring, leading to a central void. Collectively, this reasoning suggests that toroids arise from highly rigid ring-like actin bundles that fail to straighten into a set of parallel linear filaments, as would be required to transform condensates into rod-like shapes.

**Figure 3:**
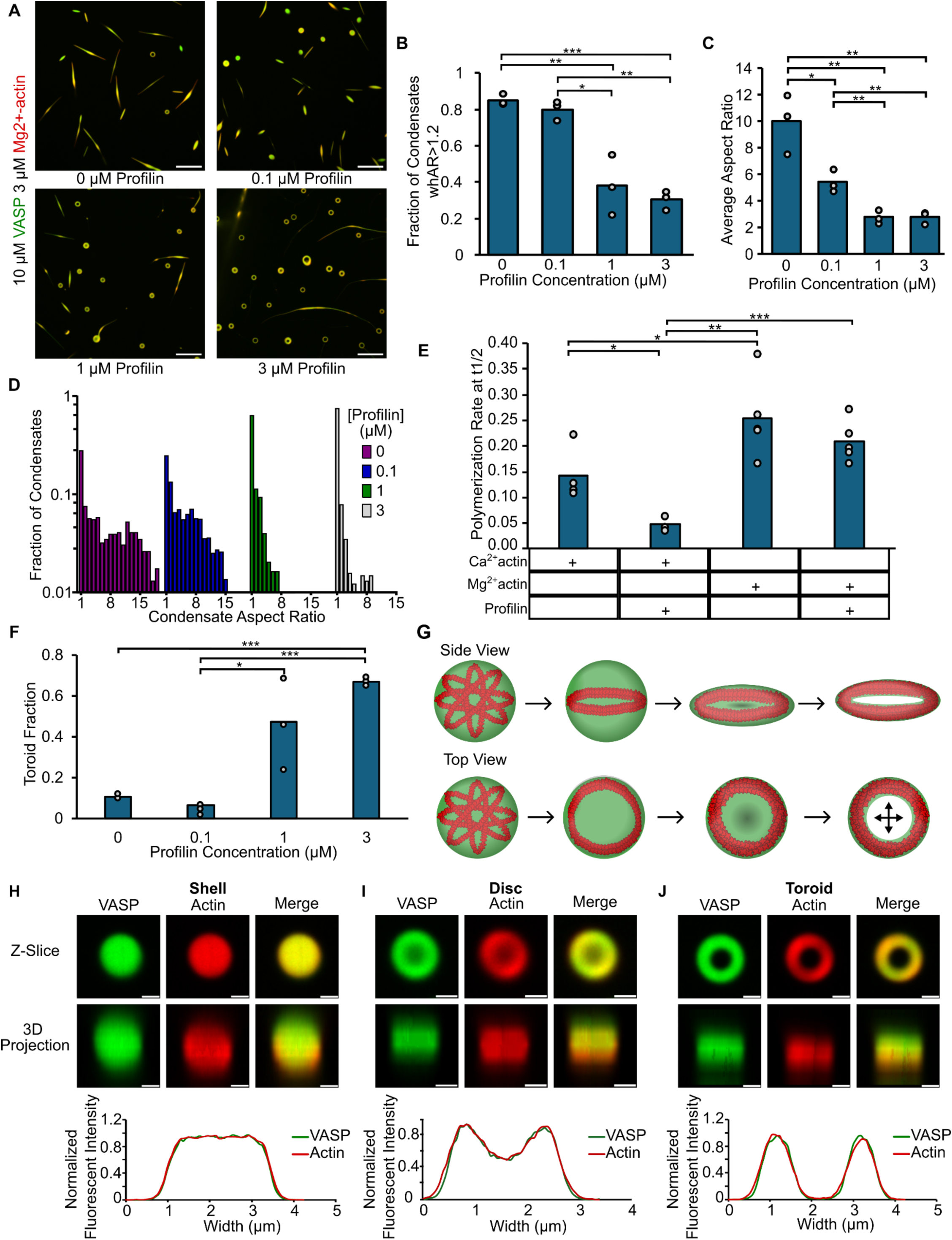
The combination of magnesium and profilin traps actin filaments and condensates in toroidal morphologies. A) Representative images showing that condensates of 10 µM VASP with 3 µM Mg^2+^-actin and increasing concentrations of profilin result in the emergence of a toroidal condensate-actin morphology. B) Quantification of the fraction of condensates with deformed morphologies, defined as having an aspect ratio greater than 1.2, for the conditions shown in A. Data are means across three independent experiments with at least 750 condensates analyzed per condition. Overlaid gray circles denote the means of each replicate. One asterisk denotes p<0.05, two asterisks denote p<.01, and three asterisks denote p<0.001 using an unpaired, two-tailed t test on the means of the individual replicates, n=3. C) Quantification of the average condensate aspect ratio for the conditions shown in A. Data are means across three independent experiments with at least 750 condensates analyzed per condition. Overlaid gray circles denote the means of each replicate. One asterisk denotes p<0.05 and two asterisks denote p<0.01 using an unpaired, two-tailed t test on the means of the individual replicates, n=3. D) Histograms showing the distribution of condensate aspect ratios for the conditions shown in A. Condensates with aspect ratios greater than 20 (corresponding to 8.8%, 0.27%, 2.15%, and 1.6% of the data, respectively, for the 0, 0.1, 1, and 3 µM conditions) were hidden to better visualize the distributions across all conditions. E) Quantification of filament assembly rates at t1/2 for pyrene actin assays of VASP condensates, Ca^2+^-actin or Mg^2+^-actin, and with or without profilin. One asterisk denotes p<.05, two asterisks denote p<.01, and three asterisks denote p<.001 using an unpaired, two-tailed t test on the calculated t1/2 values from the individual replicates, n=3. F) Quantification of the fraction of condensates that displayed a toroidal morphology for the conditions shown in A. Data are means across three independent experiments with at least 750 condensates analyzed per condition. Overlaid gray circles denote the means of each replicate. One asterisk denotes p<0.05 and three asterisks denote p<0.001 using an unpaired, two-tailed t test on the means of the individual replicates, n=3. G) Graphic depicting the proposed sphere to toroid transition with both a side view demonstrating the flattening and evacuation of the center of the protein phase. H-J) Representative images of the hypothesized progression from a sphere(H) to disc(I) to toroid(J) morphology, displaying both single slices of stacks and 3D projections. The graphs below are line profile plots along the indicated line demonstrating normalized fluorescence intensity across the condensates. Scale bars 1 μm.

Why might the combination of profilin and magnesium result in rigid ring-like bundles that do not straighten into rods? One possibility is that profilin alters the balance between filament nucleation and elongation within condensates. Specifically, profilin is known to suppress spontaneous nucleation while facilitating enhanced barbed-end elongation in the presence of elongation factors like VASP.^9,28,29,37^ Indeed, in our experimental buffer conditions, Mg²⁺-actin spontaneously polymerized at higher actin concentrations, while the addition of profilin suppressed this spontaneous filament assembly, demonstrating that it potently inhibits filament nucleation under our experimental conditions (Fig. S3).This suppression of nucleation may result in fewer total filaments within the condensates, each elongating more processively and thereby reaching a longer length. The resulting long filaments, which coil into ring-like bundles within condensates, may fail to exert sufficient asymmetric pressure on the condensate boundary to drive its deformation.

### Agent-based simulations reveal the impact of filament number and length on the deformation of condensates

To test the hypothesis that increasing filament elongation at the expense of nucleation suppresses condensate deformation, we employed an agent-based simulation of actin assembly within deformable condensates.^38,39^ Specifically, we sought to determine how the distribution of the same amount of actin in either one long filament versus a number of shorter filaments would alter condensate deformation. In all simulations, the condensate had a fixed volume, was initially spherical with a radius of 0.5 µm, and had a fixed surface tension of 2 pN/µm. We simulated four different actin configurations: 10 filaments each growing to a final length equal to one condensate circumference, one long filament growing to a length 10 times the condensate circumference, 30 filaments each growing to a length equal to one condensate circumference, and one long filament growing to 30 times the condensate circumference. VASP-actin binding kinetics were chosen to favor robust cross-linking of actin filaments by VASP.^39^ Select timepoints from these simulations are shown in **Figure 4A**. In simulations with multiple shorter filaments, the condensate aspect ratio continuously increased throughout the time course, eventually transforming the initially spherical condensate into a rod-like shape filled with parallel actin filaments. In contrast, in simulations with one long filament, the single filament lacked the rigidity to substantially deform the condensate such that it was forced to curve into a coil-like morphology, resulting in modest increases in the aspect ratios of the condensates (**Fig. 4A,B**). These condensate deformation dynamics were consistent with the evolution of the alignment angle (**Fig. 4C**). Additionally, when the VASP-actin kinetics were altered such that the filaments were not as tightly crosslinked by VASP, the single long filament was better able to uncoil and deform the condensate (**Fig. S4A**). Nevertheless, once the systems with multiple actin filaments grew to a sufficient length, actin bundles composed of many shorter filaments led to greater condensate deformation than single long filaments (**Fig. S4B,C**). These results align with recent findings that actin filaments must reach a critical length to deform condensates.^39^ The results of these simulations suggest that, if it were possible to break up long filaments, the resulting set of shorter filaments might be able to overcome the condensate surface tension, deforming condensates into rod-like geometries.

**Figure 4:**
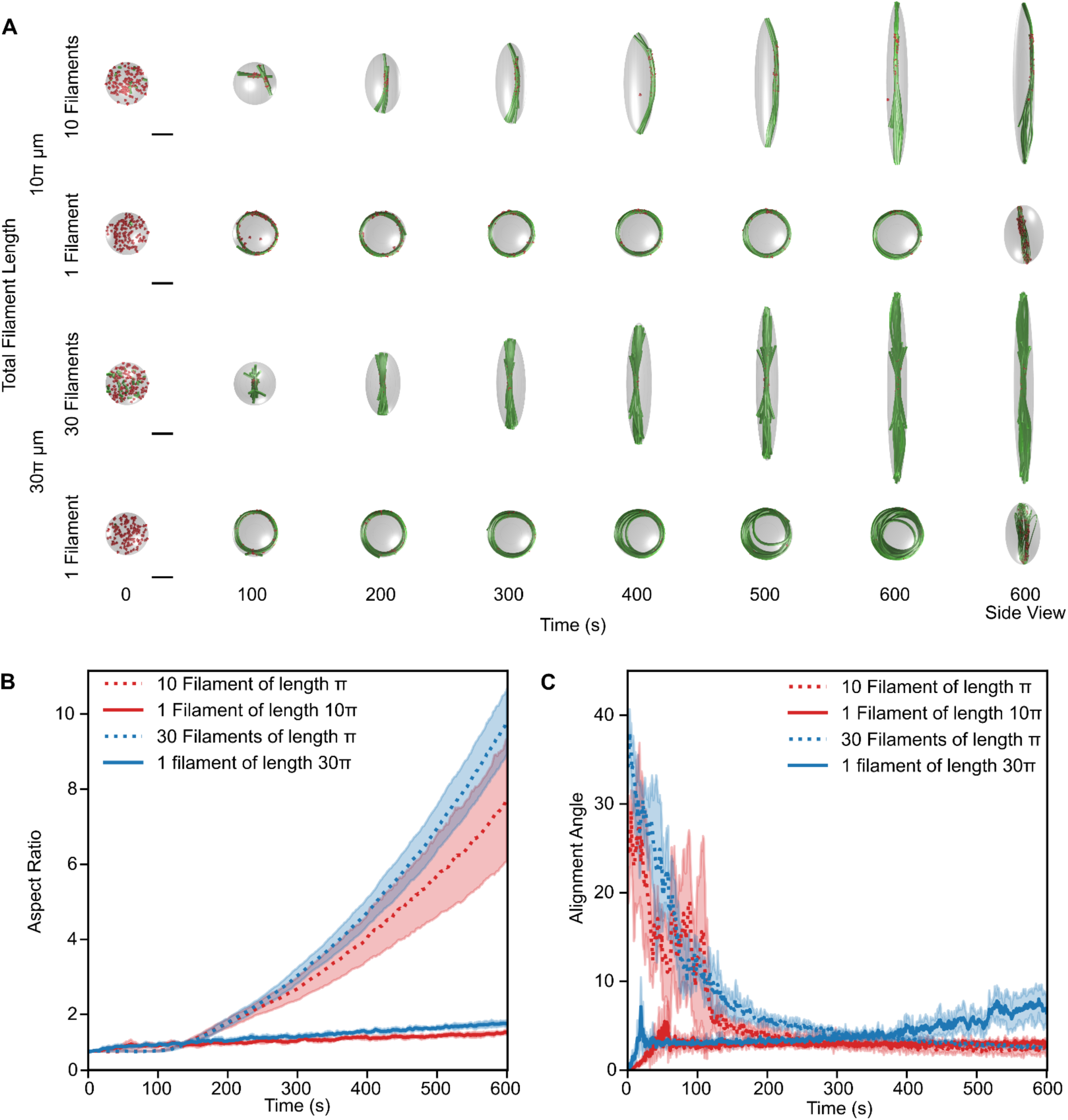
Agent-based simulations reveal the impact of filament number and length on the deformation of condensates. **A**) Representative snapshots from simulations within condensates with a deformable ellipsoidal boundary (initially spherical with R = 0.5 μm) containing a varied number of actin filaments (green) and 125 tetravalent crosslinkers (red spheres). Conditions vary the total amount of actin and the number of filaments in the simulation. The binding and unbinding rates of the tetravalent crosslinkers are fixed at kbind = 10.0 s^-1^ and kunbind = 1.0 s^-1^. The polymerization rate at the barbed end is constant and scaled for each condition such that the total filament length is reached by the end of the simulation, and neither end undergoes depolymerization. The deformable boundary has a surface tension of 2 pN/µm and an effective viscosity of 100 µm pN^-1^ s^-1^. Scale bars 0.5 µm. B) Time series showing the mean (solid line) and standard deviation (shaded area) of condensate aspect ratio for each condition. The aspect ratio is defined as the ratio between the longest and shortest axes of the ellipsoid (AR = a/c), where a ≥ b ≥ c. C) Time series showing the mean (solid line) and standard deviation (shaded area) of the alignment angle for the actin filament network in each condition. The alignment angle is defined as the angle formed by the major and minor axes of the ellipsoid that best approximates the shape of the filament network. Please refer to the Methods section for a detailed description of model development and Table S1 for the parameters used. 6 replicates were used per condition.

### Filament severing transforms toroids into rod-like actin bundles

To test the idea that rod-like morphologies could be restored by breaking up long filaments, we added the actin severing protein gelsolin to our system.^40–43^ We first added increasing concentrations of gelsolin to VASP condensates and 3 µM Ca^2+^-actin, in the absence of magnesium and profilin. Under these conditions we would expect predominantly higher aspect ratio, rod-like morphologies in the absence of gelsolin, as shown in **Fig. 1**. As the concentration of gelsolin increased, we observed a loss of condensate deformation (**Fig. 5A**). Similarly, upon addition of gelsolin, phalloidin staining revealed a lack of assembled actin structures, suggesting that gelsolin severed all long filaments in the condensate environment (**Fig. 5B**), which corresponded to a decrease in condensate aspect ratio (**Fig. 5C-E**). We next tested the impact of gelsolin in the presence of Mg^2+^-actin (3 µM) and profilin (3 µM), conditions which favored assembly of toroids (**Fig. 3A**). As the concentration of gelsolin increased (10-100 nM), we observed a pronounced transition from predominately toroidal structures to predominately rod-like structures (**Fig. 5F,G**), which corresponded with an increase in condensate aspect ratio (**Fig. 5H-J**). These findings suggest that filament severing by gelsolin generates new barbed ends that disrupt the toroid and restore rod-like architectures (**Fig. 5K**), in agreement with the predictions of our simulations (**Fig. 4**).

**Figure 5:**
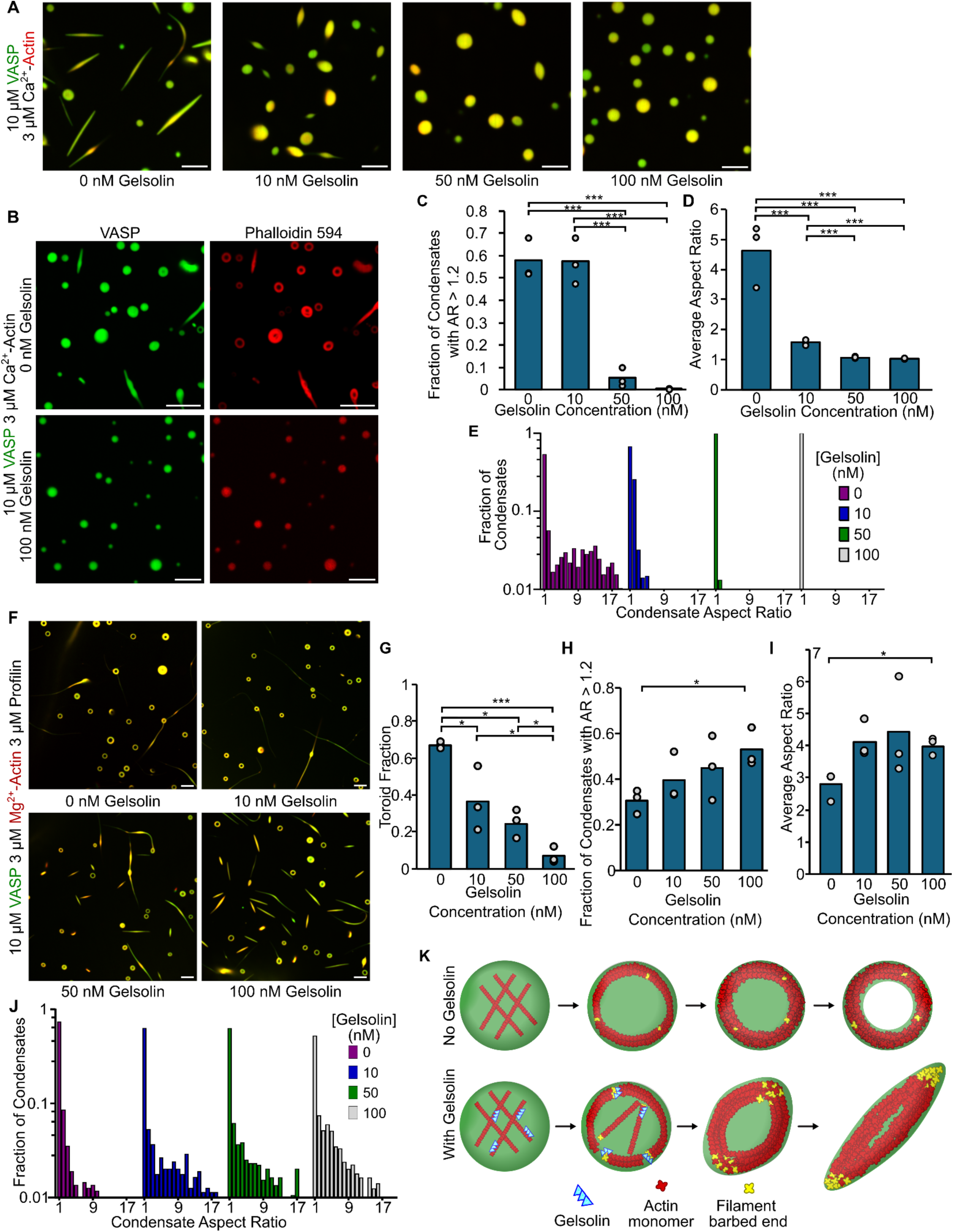
Filament severing transforms toroids into rod-like actin bundles. A) Representative confocal images showing the addition of increasing concentrations of gelsolin to 10 µM VASP condensates with 3 µM actin results in decreasing condensate deformation. B) Representative images of phalloidin staining of filamentous actin showing rings and rods of polymerized actin for conditions lacking gelsolin (Top) and the disruption of larger-scale actin filament structures with the inclusion of 100 nM gelsolin (Bottom). Conditions were 10 µM VASP condensates with 3 µM unlabeled actin in 20 mM Tris pH 7.4, 150 mM NaCl, 5 mM TCEP with 3% (w/v) PEG. C) Quantification of the fraction of condensates that have aspect ratios greater than 1.2 for the conditions shown in A. Data are means across three independent experiments with at least 750 condensates analyzed per condition. Overlaid gray circles denote the means of each replicate. Three asterisks denote p<0.001 using an unpaired, two-tailed t test on the means of the individual replicates, n=3. D) Quantification of the average condensate aspect ratio for the conditions shown in A. Data are means across three independent experiments with at least 750 condensates analyzed per condition. Overlaid gray circles denote the means of each replicate. Three asterisks denote p<0.001 using an unpaired, two-tailed t test on the means of the individual replicates, n=3. E) Histograms showing the distribution of condensate aspect ratios for the conditions shown in A. Condensates with aspect ratios greater than 20 (corresponding to 1.5% and 0.128% of the data, respectively, for the 0 and 10 nM gelsolin conditions) were hidden to better visualize the distributions across all conditions. F) The addition of increasing concentrations of gelsolin results in a transition from toroid-dominated morphologies back to rod-like condensate morphologies, suggesting that gelsolin filament severing disrupts the toroid-trapped end state. Conditions were 10 µM VASP condensates with 3 µM Mg^2+^-actin and 3 µM profilin to provide conditions that were toroid dominated based on results from Fig. 4A. G) Quantification of the fraction of condensates that display a toroidal morphology for the conditions shown in E. Data are means across three independent experiments with at least 800 condensates analyzed per condition. Overlaid gray circles denote the means of each replicate. One asterisk denote p<0.05 and three asterisks denote p<.001 using an unpaired, two-tailed t test on the means of the individual replicates, n=3. H) Quantification of the fraction of condensates that have aspect ratios greater than 1.2 for the conditions shown in F. Data are means across three independent experiments with at least 750 condensates analyzed per condition. Overlaid gray circles denote the means of each replicate. One asterisk denotes p<0.05 using an unpaired, two-tailed t test on the means of the individual replicates, n=3. I) Quantification of the average condensate aspect ratio for the conditions shown in F. Data are means across three independent experiments with at least 750 condensates analyzed per condition. Overlaid gray circles denote the means of each replicate. One asterisk denotes p<0.05 using an unpaired, two-tailed t test on the means of the individual replicates, n=3. J) Histograms showing the distribution of condensate aspect ratios for the conditions shown in F. Condensates with aspect ratios greater than 20 (corresponding to 1.3%, 1.9%, 2.6%, and 0.59% of the data, respectively, for the 0, 10, 50, and 100 nM gelsolin conditions) were hidden to better visualize the distributions across all conditions. K) Cartoon depicting how gelsolin-mediated filament severing inhibits trapped ring and toroid formation, and results in a return to rod-like condensate morphologies.

## Discussion

Here, we show that deformation of protein condensates by actin filaments requires a balance between filament nucleation and elongation. When filament elongation dominates over nucleation, long actin filaments fail to apply enough asymmetric pressure on condensate boundaries to deform them into rod-like geometries. Specifically, when magnesium, which favors filament growth, and profilin, which inhibits filament nucleation, were added simultaneously to VASP condensates, ring-like bundles of long actin filaments deformed initially spherical condensates into disc and toroid-like morphologies that failed to proceed to linear, rod-like geometries, in agreement with agent-based simulations. The addition of intermediate concentrations of the filament-severing protein, gelsolin, to these structures created free barbed ends that grew to form rod-like morphologies. However, in the presence of an excess of gelsolin, filament severing dominated over filament elongation, resulting in a population of very short filaments that failed to deform condensates. Taken together, the results of our experiments illustrate that both filament elongation and nucleation are required for deformation of protein condensates into high aspect ratio structures by growing actin filaments.

Computational modeling of actin networks in droplets has emerged as a powerful tool to generate experimentally testable hypotheses. Our previous modeling work showed that kinetic trapping is an important mechanism underlying the formation of ring-like bundles in droplets,^19^ and that mechanochemical feedback between the droplet properties and the actin network organization determines the deformation dynamics.^39^ Here, our simulations suggest that short, linear bundles of actin are more effective at exerting forces on the droplet interface leading to rod-like structures. In contrast, a single long filament, which simulations predict will be forced by the droplet surface tension to form a coil, is not expected to generate sufficient asymmetric pressure to deform the droplet into a high aspect ratio structure.

Taken together, our experimental and computational results demonstrate that the kinetics of F-actin nucleation and elongation ultimately control condensate morphology. By modulating the number and length of filaments within confined, deformable environments, phase-separated protein assemblies direct the structural outcome of actin organization, producing distinct morphologies such as rings, toroids, or rods. It has long been understood that proteins which cross link actin networks by binding to multiple actin filaments can determine the architecture of the actin network.^44^ Coupling this process to the surface tension of the condensate interface results in a variety of higher order morphologies. As an increasing number of actin-interacting proteins are being shown to undergo phase separation,^13–21^ understanding how their combined activities influence filament distribution and network morphologies will be essential for understanding the mechanisms that govern actin remodeling in complex cellular contexts.

## Methods

Reagents Tris base, NaCl, EGTA, Tris(2-carboxyethyl)phosphine (TCEP), poly-L-lysine, and Atto 594 maleimide were purchased from Sigma-Aldrich. Atto 488 maleimide was purchased from Thermo Fisher Scientific. Phalloidin-iFluor594 was purchased from Abcam. Amine-reactive PEG (mPEG-succinimidyl valerate, MW 5,000) was purchased from Laysan Bio. Rabbit muscle actin, gelsolin, and profilin were purchased from Cytoskeleton.

### Plasmids

A pET vector encoding the ‘cysteine light’ variant of human VASP (pET-6xHis-TEV-KCK-VASP(CCC-SSA)) was a gift from Scott Hansen.

### Protein Purification

The pET-His-KCK-VASP(CCC-SSA) plasmid was transformed into Escherichia coli BL21(DE3) competent cells (NEB, cat. no. C2527). Cells were grown at 30 °C to an optical density (OD) of 0.8. Protein expression was performed as described previously with some modifications as follows.^18^ Expression of VASP was induced with 0.5 mM isopropylthiogalactoside (IPTG), and cells were shaken at 200 rpm at 12 °C for 24 h. The rest of the protocol was carried out at 4 °C. Cells were pelleted from 2 L cultures by centrifugation at 4,785g (5,000 rpm in Beckman JLA-8.100) for 20 min. Cells were resuspended in 100 mL lysis buffer (50 mM sodium phosphate pH 8.0, 300 mM NaCl, 5% glycerol, 0.5 mM TCEP, 10 mM imidazole, 1 mM phenylmethyl sulphonyl fluoride (PMSF)) plus EDTA-free protease inhibitor tablets (1 tablet per 50 mL, Roche, cat. no. 05056489001), 0.5% Triton-X100, followed by homogenization with a dounce homogenizer and sonication (4 × 2,000 J). The lysate was clarified by ultracentrifugation at 125,171g (40,000 rpm in Beckman Ti45) for 30 min. The clarified lysate was then applied to a 10 mL bed volume Nickel nitrilotriacetic acid (Ni-NTA) agarose (Qiagen, cat. no. 30230) column, washed with 10 column volumes of lysis buffer plus EDTA-free protease inhibitor tablets (1 tablet per 50 mL), 20 mM imidazole, 0.2% Triton X-100, followed by washing with 5×CV of lysis buffer plus 20 mM imidazole. The protein was eluted with elution buffer (50 mM Tris, pH 8.0, 300 mM NaCl, 5% glycerol, 250 mM imidazole, 0.5 mM TECP, EDTA-free protease inhibitor tablets (1 tablet per 50 mL)). The protein was further purified by size exclusion chromatography with Superose 6 resin. The resulting purified KCK-VASP was eluted in storage buffer (25 mM HEPES pH 7.5, 200 mM NaCl, 5% glycerol, 1 mM EDTA, 5 mM DTT). Single-use aliquots were flash-frozen using liquid nitrogen and stored at −80 °C until the day of the experiment.

### Protein labeling

The VASP construct used in these studies is a previously published ‘cysteine light’ mutant that replaced the three endogenous cysteines with two serines and an alanine. A single cysteine was then introduced at the N-terminus of the protein to allow selective labeling with maleimide dyes. This mutant was found to function in an indistinguishable manner from the wild-type proteins.^28^ Thus, VASP was labeled at the N-terminal cysteine using maleimide-conjugated dyes. VASP was buffer exchanged into 20 mM Tris (pH 7.4), 150 mM NaCl buffer to remove DTT from the storage buffer, and then incubated with dye for two hours at room temperature. Free dye was then removed by applying the labeling reaction to a spin column packed with Sephadex G-50 Fine DNA Grade (GE Healthcare GE17-0573-01) hydrated with buffer containing 20 mM Tris pH 7.4, 150 mM NaCl, and 5 mM TCEP.

Monomeric actin was labeled using maleimide-conjugated dyes. Dyes were incubated with G-actin at a 2-fold molar excess for 2 hours at room temperature before being separated from the labeled protein by applying the labeling reaction to a spin column packed with Sephadex G-50 Fine DNA Grade (GE Healthcare GE17-0573-01) hydrated with A buffer (5 mM Tris-HCL (pH 8), 0.2 mM ATP and 0.5 mM DTT pH 8). The labeled protein was then centrifuged at 100,000 x G for 10 min at 4 degrees Celsius to remove aggregates before being flash-frozen in single-use aliquots.

Profilin was labeled using NHS ester dyes. Dye was incubated with profilin at a 1:1 molar ratio for 30- 45 minutes on ice before being separated from labeled protein before being run through a Zeba Biotin and Dye removal column (7k MWCO) equilibrated with 20 mM Tris pH 7.4, 150 mM NaCl, and 5 mM TCEP.

### Protein condensate formation and actin filament assembly

Condensates composed of VASP were formed by mixing the given concentration of protein (see text) with 3% (w/v) PEG 8000 in 20 mM Tris pH 7.4, 5 mM TCEP, 200 µM ATP, and 150 mM NaCl. PEG was added last to induce condensate formation after the protein was evenly dispersed in the solution. For conditions facilitating Mg^2+^ exchange, 1 mM MgCl2 and 1 mM EGTA were included in the buffer. Gelsolin was added to the protein mix prior to PEG addition for condensates consisting of VASP and gelsolin. All protein concentrations listed are the monomeric concentrations.

For actin assembly assays within condensates, condensates were formed for ten minutes (with time starting after PEG addition), and then G-actin was added to the condensate solution and allowed to assemble for 15 minutes before imaging. For phalloidin-actin assays, unlabelled G-actin was added to pre-formed protein condensates and allowed to assemble for 10 min. Phalloidin-iFluor594 was then added to stain filamentous actin for 10 min before imaging. For assays that included profilin, the corresponding amounts of actin and profilin were mixed and allowed to incubate for 15 minutes prior to addition to the condensate mix in order to facilitate profilin-actin formation.

For FRAP experiments, condensates formed from the various proteins were observed in solution under the conditions given in the text. A region within the condensates was bleached, and consecutive images were taken every three seconds to monitor fluorescence recovery over time.

### Microscopy

Samples were prepared for microscopy in 3.5mm diameter wells formed using biopsy punches to create holes in 1.6 mm thick silicone gaskets (Grace Biolabs) on Hellmanex III cleaned, no. 1.5 glass coverslips (VWR). Coverslips were passivated using poly-L-lysine conjugated PEG chains (PLL-PEG). To prevent evaporation during imaging, an additional small coverslip was placed on top of the gasket to seal the well. Fluorescence microscopy was done using an Olympus SpinSR10 spinning disk confocal microscope with a Hamamatsu Orca Flash 4.0V3 Scientific CMOS camera. FRAP was done using the Olympus FRAP unit 405 nm laser.

PLL-PEG was prepared as described previously with minor modifications as follows.^45^ Briefly, amine-reactive mPEG succinimidyl valerate was conjugated to poly-L-lysine at a molar ratio of 1:5 PEG to PLL. The conjugation reaction takes place in 50 mM sodium tetraborate solution, pH 8.5, and is allowed to react overnight at room temperature while continuously stirring. The final product is then buffer exchanged to PBS pH 7.4 using 7000 MWCO Zeba spin desalting columns (Thermo Fisher) and stored at 4 °C.

### Pyrene Assay

Pyrene-labeled actin was obtained from Cytoskeleton Inc. (AP05-A). Lyophilized labeled actin was resuspended to 465 μM with cold distilled water and stored per the manufacturer’s instructions. For polymerization, a stock solution of 465 μM pyrene-labeled G-actin was diluted with fresh G-Buffer (2 mM Tris-HCl, 0.5 mM DTT, 0.2 mM CaCl2, 0.2 mM ATP) to a working concentration of 10 μM with G-Buffer. A 6 μM 10% pyrene-labeled actin mix was made through dilution with unlabeled G-actin. The G-actin solution was left on ice to depolymerize for 1 h before ultracentrifugation at 7300×g for 30 min at 4 °C, after which the supernatant was then collected. For the varying protein conditions, 50 μL of protein mix was added to the wells of a black flat-bottom 96-well assay plate (Corning #3915) and mixed gently before immediate transfer to the plate reader. For conditions with actin alone, the protein mix was just experimental buffer (20 mM Tris pH 7.4, 150 mM NaCl, 5 mM TCEP, 200 μM ATP). For conditions with Mg^2+^ actin 1 mM MgCl2 and 1 mM EGTA were included to facilitate divalent cation exchange. For conditions with VASP, the protein mix consisted of 20 μM VASP. For conditions with PEG, PEG was added for a final concentration of 3% (w/v) to initiate condensate formation. For conditions with profilin, profilin was added to the actin mix for a final concentration of 3 μM. Protein fluorescence measurements were then taken every 30 seconds for 3 minutes at excitation and emission wavelengths of 360 ± 20 nm and 405 ± 10 nm, respectively, with a plate reader to establish a fluorescence baseline. 50 μL of the G-actin mix was then pipetted into each well of the 96-well assay plate and fluorescence measurements were then taken every 30 seconds for 2 hours. t1/2, actin polymerization rate at t1/2 and initial nucleation rates were calculated per ^46^. For nucleation rate calculations, fluorescence intensity values up to 2 or 2.5 seconds after addition of G-actin to the wells were used.

### Image Analysis

Image J was used to quantify the distribution of condensate characteristics. Specifically, condensates were selected using thresholding in the brightest channel and shape descriptors (i.e. diameter, aspect ratio, etc.), and protein fluorescent intensities were measured using the built-in analyze particles function. For aspect ratio analysis, condensates that had come into contact with other condensates were removed from the analysis to avoid any skewing of data from misrepresentation of single condensate deformation.

FRAP data were analyzed using ImageJ, where fluorescence recovery over time was measured and then normalized to the maximum pre-bleach intensity. Recovery was measured for condensates of similar diameters and photobleached region size.

Partitioning data was calculated using the average intensities of the condensed protein phase and the bulk solution, with partitioning defined as the ratio of the intensity inside the condensate to outside the condensate. Images were cropped so that only condensates from the middle ninth of the field of view were analyzed to avoid any error from potential non-uniform illumination across the imaging field.

### Modeling

#### Model Development

Simulations were performed in Cytosim (https://gitlab.com/f-nedelec/cytosim), an agent-based modeling framework that simulates the chemical dynamics and mechanical properties of cytoskeletal filament networks.^47^ Filament dynamics and diffusing species are modeled by numerically solving a constrained Langevin framework in a viscous medium, a stochastic differential equation for describing Brownian motion, to resolve position evolution at short time intervals. Each actin filament is represented by a series of segments, each with a maximum length Lseg = 0.1 µm. Filaments elongate deterministically at a constant rate kgrow such that the filaments grow to the desired maximum length by the end of the simulation at 600 s. Tetrameric VASP crosslinkers are modeled as spheres with four actin-binding sites with actin-binding kinetics governed by rate parameters kbind and kunbind. Each binding site has a specified binding radius that determines the spherical volume within which binding partners are considered as part of the binding reaction. Unbinding reactions are governed by a Bell’s law model representation of slip bond unbinding kinetics where the rate constant is given by the unbinding rate.^48,49^ Steric repulsion potentials between diffusing crosslinkers and filaments are employed to avoid spatial overlap of species.

The dynamically deformable ellipsoid framework for Cytosim was developed in as a computationally expedient method for investigating the role of cortical tension in microtubule assembly within red blood cells.^38^ We incorporate a deformable ellipsoid geometry that dynamically adjusts the space and axes with each time step to model the condensate surface as a simple continuously deformable surface constrained by a constant volume.^39^ Deformation forces are calculated as a force balance between the forces generated by the confined filaments, the resistance to deformation governed by the interfacial surface tension parameter σsurface, and the pressure which serves as a Lagrange multiplier calculated to enforce volume incompressibility.^38^ As the simulation evolves with time, the force balance is projected along the three axes of the ellipsoid and the speed of deformation along these axes is calculated and attenuated by a damping parameter given by the effective viscosity, µeffective, which slows the deformation without affecting the final droplet shape.^38^ A full formulation of this model can be found in Dmitrieff et al. 2017.^38^

Detailed methods and parameters for the modeling component of this work can be found in **Table S1**. **Quantification and Statistical Analysis**

Statistical details of experiments can be found in the corresponding figure captions, including replicate numbers, *n* values, significance tests used, and significance thresholds.

### Resource Availability Lead contact

Further information and requests for resources and reagents should be directed to and will be fulfilled by the lead contact, Jeanne Stachowiak.

### Materials Availability

All unique/stable reagents generated in this study are available from the lead contact with a completed material transfer agreement.

### Data and code availability

Microscopy data and analysis will be shared by the lead contact upon request. The custom Cytosim code and input files used to generate trajectories and the python scripts used to analyze trajectories are available upon request from the lead contact upon request. Any additional information required to reanalyze the data reported in this work paper is available from the lead contact upon request.

## Acknowledgments

This research was primarily supported by the National Science Foundation through the Center for Dynamics and Control of Materials: an NSF MRSEC under Cooperative Agreement No. DMR- 2308817. Additionally, this research was supported by grants from the NIH to J.C.S. (R35GM139531) and by the NSF through a Modulus Grant MCB 2327244 to P.R. and J.C.S. D.J. was supported by the Ronald E. McNair Scholars Program.

## Author Contributions

C.W., U.E., B.B., J.A., P.R., and J.C.S. designed experiments. C.W., U.E., B.B., J.A., P.R., and J.C.S. wrote and edited the manuscript. C.W., U.E., D.J., P.R., and J.C.S. performed experiments and analyzed data. All authors consulted on manuscript preparation and editing.

## Declaration of Interests

P.R. is a consultant for Simula Research Laboratories in Oslo, Norway, and receives income. The terms of this arrangement have been reviewed and approved by the University of California, San Diego in accordance with its conflict-of-interest policies.

## Supplementary Figures

**Figure S1:**
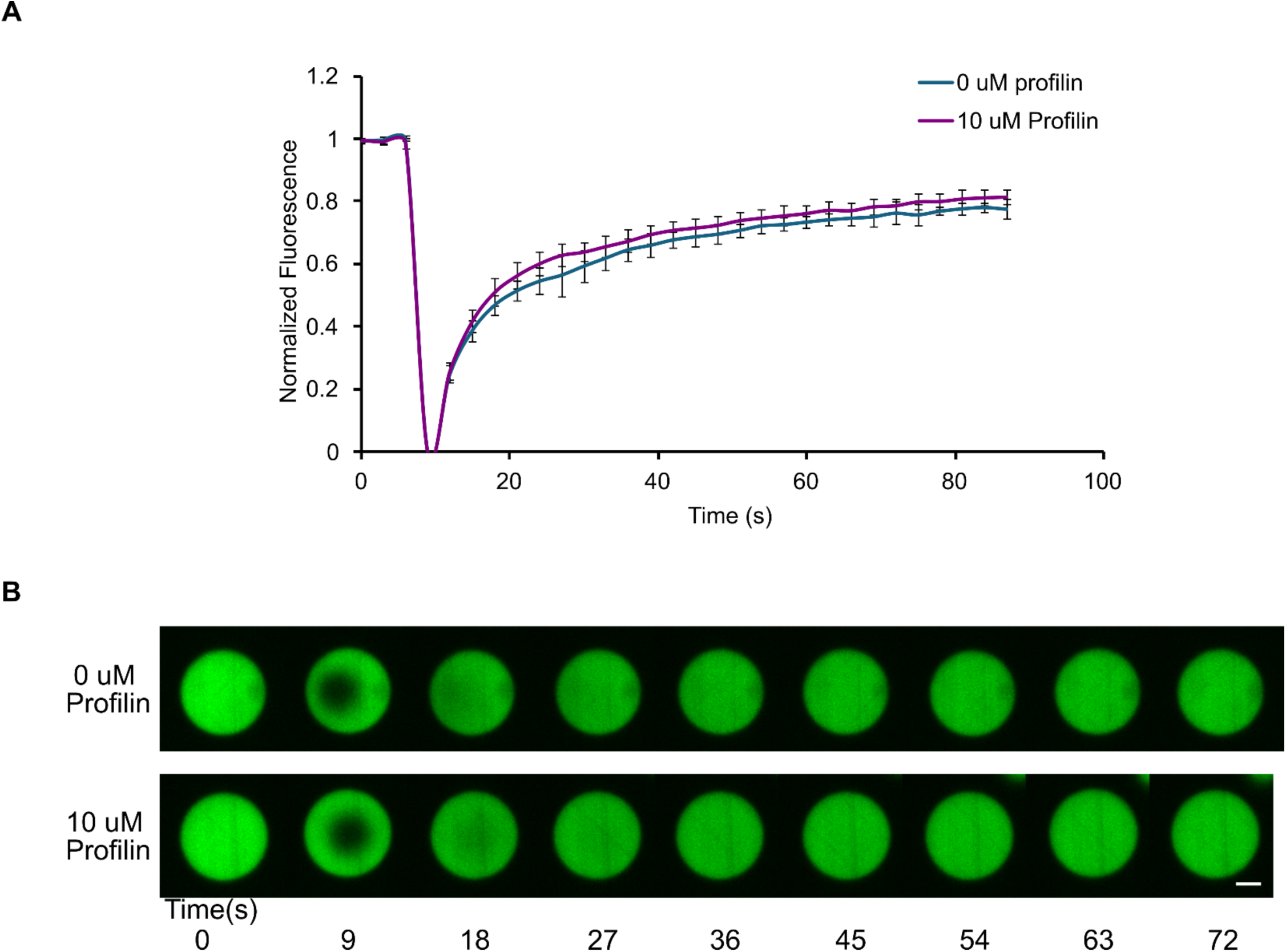
FRAP recovery of VASP condensates with and without Profilin. **A)** Plot of average fluorescence recovery +- SD after photobleaching for VASP condensates across n = 9 independent samples for conditions with and without added profilin. **B)** Representative images of fluorescent recovery after photobleaching of VASP condensates in conditions with and without added profilin. Scale bars 2 µm.

**Figure S2:**
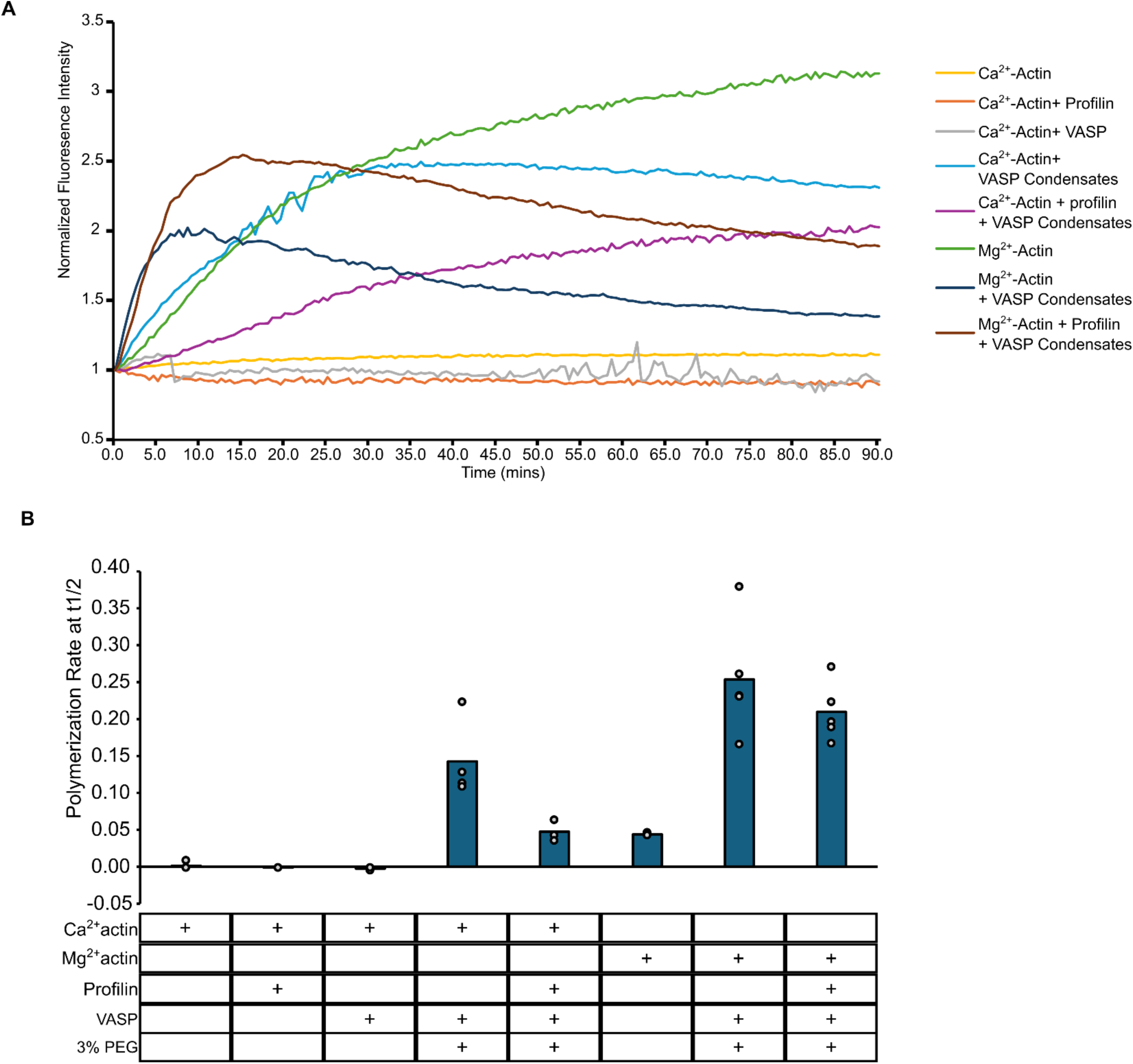
Full Pyrene Assay Data: **A)** Pyrene assay fluorescence intensity curves for actin polymerization assays. Values have been normalized such that the starting intensity is equal to 1 and increases represent a x-fold increase in pyrene fluorescence. Curves represent the average of at least three individual replicates. **B)** Quantification of the actin polymerization rate ± SD at t1/2 for the conditions shown in **A.** Data are means across at least three replicates. Overlaid gray circles denote the means of each replicate. The table underneath notes the components that were included in the assay. The inclusion of 3% PEG denotes conditions in which VASP condensates were formed.

**Figure S3:**
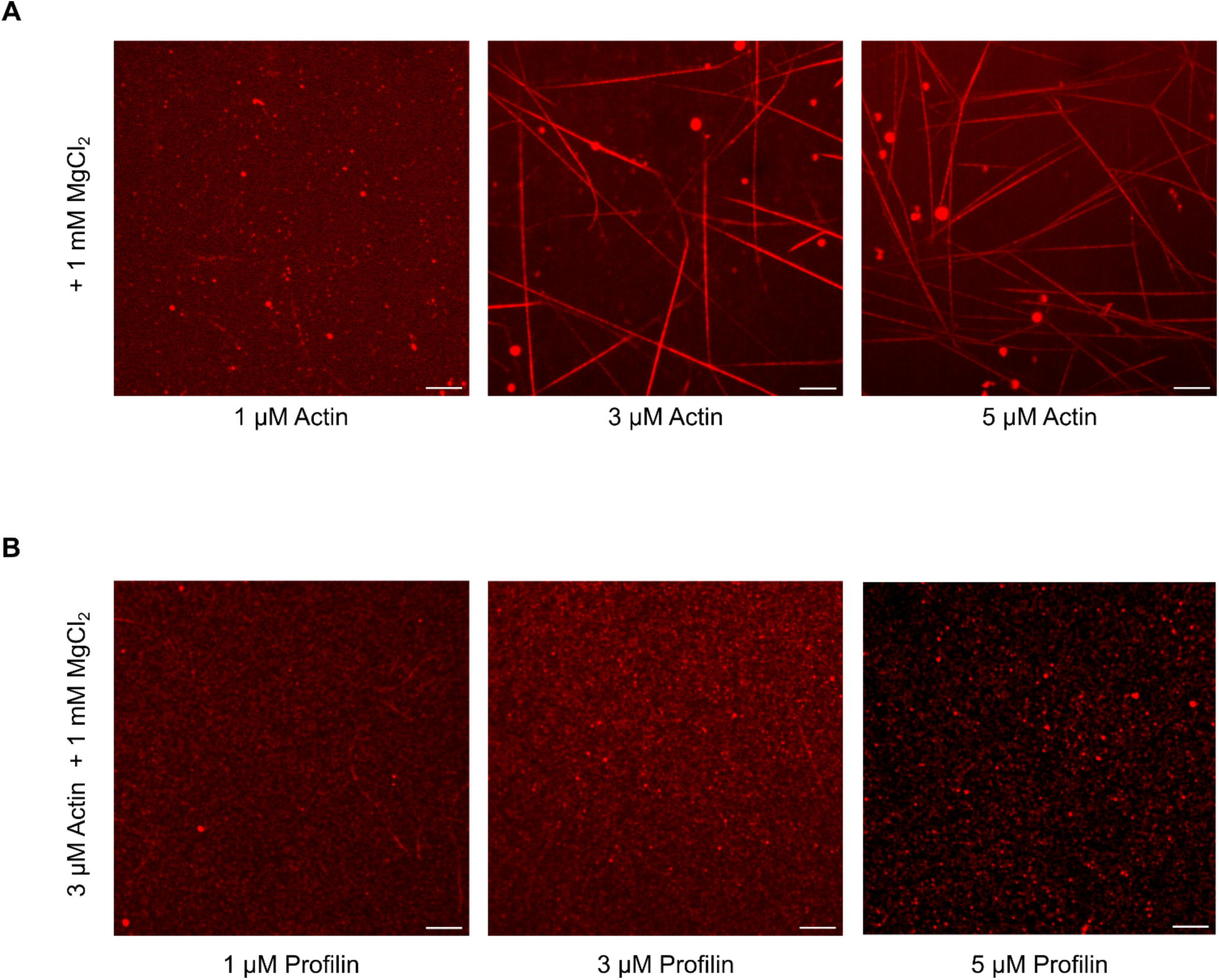
Mg^2+^ and profilin affect spontaneous actin assembly in experimental conditions: **A)** With buffer conditions facilitating Mg^2+^ exchange (1 mM added MgCl2 and EGTA), actin spontaneously polymerizes at higher concentrations in the experimental setup used for condensate experiments. **B)** At conditions that facilitated spontaneous filament assembly in **A** (3 µM Actin with 1 mM added MgCl2 and EGTA), the addition of profilin in increasing concentrations suppressed spontaneous actin filament assembly in the experimental setup used for condensate experiments. Scale bars 5 µm.

**Figure S4:**
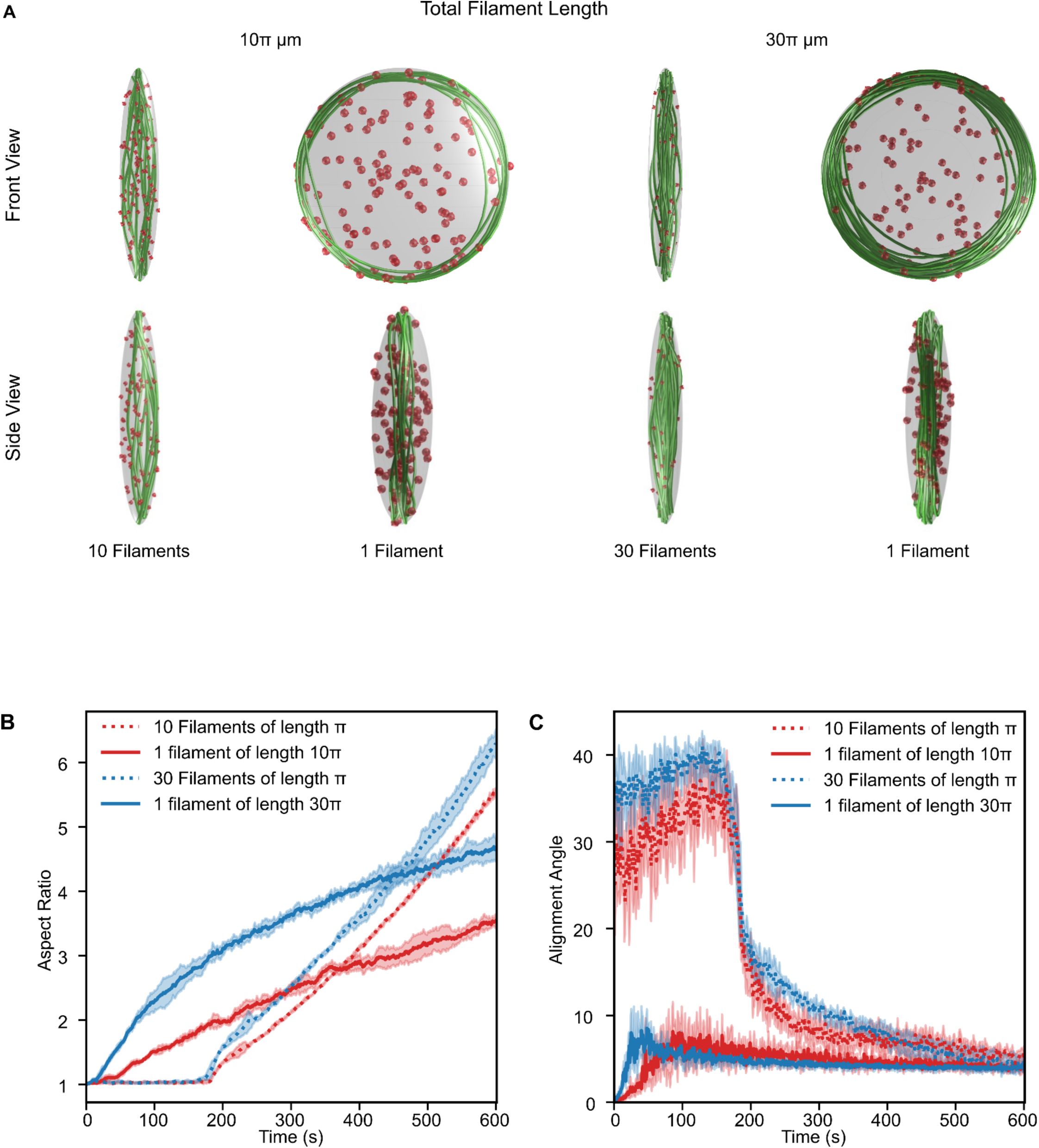
Agent-based simulations reveal the impact of filament number and length on the deformation of condensates in a weaker crosslinking environment. **A)** Representative final snapshots (t = 600 s) of both the front view (top) and a side view (bottom) from simulations within condensates with a deformable ellipsoidal boundary (initially spherical with R = 0.5 μm) containing a varied number of actin filaments (green) and 125 tetravalent crosslinkers (red spheres). Conditions vary the total amount of actin and the number of filaments in the simulation. The binding and unbinding rates of the tetravalent crosslinkers are fixed at kbind = 0.1 s^-1^ and kunbind = 1.0 s^-1^. The polymerization rate at the barbed end is constant and scaled for each condition such that the total filament length is reached by the end of the simulation, and neither end undergoes depolymerization. The deformable boundary has a surface tension of 2 pN/µm and an effective viscosity of 100 µm pN^-1^ s^-1^. **B)** Time series showing the mean (solid line) and standard deviation (shaded area) of condensate aspect ratio for each condition. The aspect ratio is defined as the ratio between the longest and shortest axes of the ellipsoid (AR = a/c) where a ≥ b ≥ c. **C)** Time series showing the mean (solid line) and standard deviation (shaded area) of the alignment angle for the actin filament network in each condition. The alignment angle is defined as the angle formed by the major and minor axes of the ellipsoid that best approximates the shape of the filament network. Please refer to the **Methods** section for a detailed description of the model and **Table S1** for the parameters used. 6 replicates were used per condition.

## Supplementary Tables

**Table S1:** Table of parameters used in the Cytosim model.

| Parameter | Value | Notes/Reference |
| --- | --- | --- |
| Total time ($T_{sim}$) | 600 s | |
| Time step | 0.002 s |  |
| Condensate viscosity | 0.5 pN s/μm<sup>2</sup> | 500x water;<br>Chosen based on protein condensate viscosities.<sup>50</sup> |
| **Boundary** |  |  |
| Initial Shape | Spherical, constrained to a deformable ellipsoid. |  |
| Surface Tension $\sigma_{surface}$ | 2 pN/μm | Chosen to ensure the observation of ring and rod structures within the kinetic parameters used as previously determined.<sup>39</sup> |
| Effective Viscosity: $\mu_{effective}$ | 100 pN s/μm | |
| Radius | 0.5 μm |  |
| Boundary repulsion stiffness | 200 pN/μm for actin filaments;<br>100 pN/μm for crosslinking molecules | This specifies the spring stiffness that acts on the discretized points of each actin filament and crosslinking molecule if the point lies outside the specified condensate boundary. The force on each point is dependent on the distance that it lies beyond the condensate boundary. |
| **Actin filaments** |  |  |
| Segmentation length $L_{seg}$ | 0.1 μm (100 nm) | |
| Maximum length | 10 Filaments: $\pi$ μm<br>10x Length Filament: $10\pi$ μm<br>30 Filaments: $\pi$ μm<br>30x Length Filament: $30\pi$ μm | Varied in this study. |
| Polymerization rate $k_{\text{grow}}$ | 10 Filaments: 0.00506933 $\mu\text{m/s}$<br>10x Length Filament: 0.0521933 $\mu\text{m/s}$<br>30 Filaments: 0.00506933 $\mu\text{m/s}$<br>30x Length Filament: 0.156913 $\mu\text{m/s}$ | Only barbed end elongation occurs at a rate that is calculated such that the desired final filament length is reached at 600 s. |
| Brownian ratchet force for polymerization | 10 pN | 51 |
| Actin flexural rigidity $k_{\text{bend}}$ | 0.075 pN$\cdot\mu\text{m}^2$ | 52 |
| Actin steric repulsion $k_{\text{steric}}$ | Radius 3.5 nm<br>Stiffness 1.0 pN/$\mu\text{m}$ | Chosen to ensure the observation of ring structures within the kinetic parameters used as previously determined.<sup>19</sup> |
| **Tetravalent crosslinkers (VASP)** |  |  |
| Radius | 30 nm |  |
| Diffusion rate | 10 $\mu\text{m}^2/\text{s}$ | |
| Concentration of tetramers | 0.40 $\mu\text{M}$ [125 tetramers when the initial condensate radius = 0.5 $\mu\text{m}$] | |
| Actin-binding rate (Ring Kinetics) | 10.0 (1/s) | Kinetic parameter chosen to ensure the observation of ring structures as previously determined.<sup>39</sup> |
| Actin-binding rate (Rod Kinetics) | 0.1 (1/s) | Kinetic parameter chosen to ensure the observation of rod structures as previously determined.<sup>39</sup> |
| Actin-binding distance | 30 nm |  |
| Actin-binding valency | 4 | Each spherical molecule approximates a VASP tetramer. |
| Zero-force actin-unbinding rate (Ring and Rod Kinetics) | 1.0 (1/s) | Kinetic parameter chosen to ensure the observation of ring and rod structures as previously determined.<sup>39</sup> |
| Actin-unbinding force | 10 pN | Typical values for passive crosslinkers.<sup>53</sup> |
| VASP steric repulsion | Radius 30 nm<br>Stiffness 10 pN/$\mu\text{m}$ | Chosen to ensure the observation of ring structures within the kinetic parameters used in this study as determined from a previous study on tetramers.<sup>19</sup> |

## Notes

### Competing Interest Statement

The authors have declared no competing interest.

